# Single and binary protein electroultrafiltration using poly(vinyl-alcohol)-carbon nanotube (PVA-CNT) composite membranes

**DOI:** 10.1101/2020.01.29.924753

**Authors:** Raymond Yeung, Xiaobo Zhu, Terence Gee, Ben Gheen, David Jassby, Victor G. J. Rodgers

**Affiliations:** Department of Bioengineering, University of California, Riverside, Riverside, California, United States of America; Department of Civil and Environmental Engineering, University of California, Los Angeles, Los Angeles, California, United States of America

## Abstract

Electrically conductive composite ultrafiltration membranes composed of carbon nanotubes have exhibited efficient fouling inhibition in wastewater treatment applications. In the current study, poly(vinyl-alcohol)-carbon nanotube membranes were applied to fed batch crossflow electroultrafiltration of dilute (0.1 g/L of each species) single and binary protein solutions of α-lactalbumin and hen egg-white lysozyme at pH 7.4, 4 mM ionic strength, and 1 psi. Electroultrafiltration using the poly(vinyl-alcohol)-carbon nanotube composite membranes yielded temporary enhancements in sieving for single protein filtration and in selectivity for binary protein separation compared to ultrafiltration using the unmodified PS-35 membranes. Assessment of membrane fouling based on permeate flux, zeta potential measurements, and scanning electron microscopy visualization of the conditioned membranes indicated significant resulting protein adsorption and aggregation which limited the duration of improvement during electroultrafiltration with an applied cathodic potential of −4.6 V (vs. Ag/AgCl). These results imply that appropriate optimization of electroultrafiltration using carbon nanotube-deposited polymeric membranes may provide substantial short-term improvements in binary protein separations.

## Introduction

Carbon nanotubes (CNT) have drawn significant attention as an ideal conductive material to modify UF membranes [1, 2]. Composite UF membranes combine the properties of good electrical conductivity, robust mechanical strength, and efficient filtration performance. Electrically conducting membranes with tunable charged surfaces under an applied electrical potential take advantage of electrophoretic and electrostatic contributions to the transport of charged particles (Fig 1). Extensive work has been performed to develop and investigate the use of electrically conductive composite CNT-polymer composite UF membranes for water treatment [3–8].

**Fig 1.**
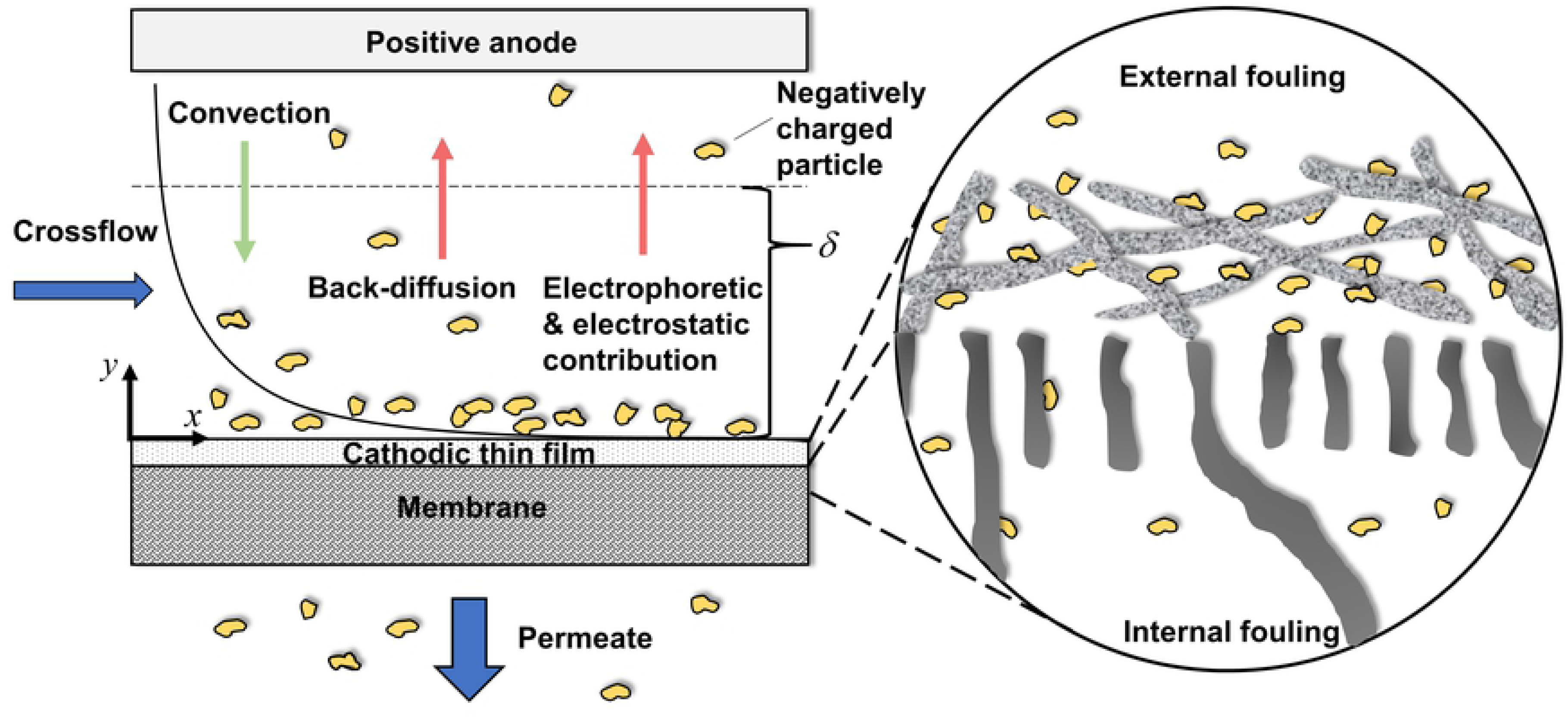
Schematic view of crossflow electroultrafiltration with an electrically conductive, cathodic membrane. As can be seen, unlike electroultrafiltration processes with electrodes on each side of the membrane, the external electric field terminates with the poly(vinyl-alcohol)-carbon nanotube layer, which is upstream of the semipermeable, polymeric membrane.

Jassby et al. [5-7, 9-11] developed methods of coating commercial polymeric UF membranes with a thin layer of a poly(vinyl-alcohol) (PVA) polymer cross-linked with multiwalled carbon nanotubes to form highly electrically conductive PVA-CNT composite membranes. In a study of the effect of moderate applied electric potentials (−1.5 and −3.4 V vs Ag/AgCl references) on fouling of high concentrations (3-5 g/L) of synthetic wastewater containing negatively charged alginic acid during EUF using PVA-CNT composite membranes, Dudchenko and Jassby et al. observed substantial fouling inhibition and reduction of operating pressure in a constant flux system [6]. Moreover, Ronen and Jassby et al. used PVA-CNT composite membranes for treatment of aqueous solutions containing bacteria and reported significant reduction in bacterial attachment upon applying both anodic (1.2 V vs. Ag/AgCl) and cathodic potentials (−0.6 V vs. Ag/AgCl) [7].

While electroultrafiltration using composite conductive CNT-polymer UF membranes that also act as polarizable electrodes have been used extensively for processing solutions containing charged molecules, the method has yet to be evaluated for its potential for enhancement in protein filtration and separation. Several investigators have shown that electrostatic interactions between charged proteins and charged UF membranes strongly affect protein sieving [12–16]. However, there have been no fundamental studies of the electrostatic effects from an externally applied electrical potential on protein transport during EUF with a CNT-based composite membrane. Electrostatic interactions, hydrophobic interactions, hydrogen bonding, and π-π interactions are reported to play key roles in CNT/protein binding. Although there are studies suggesting π-π interactions stacking interactions as the main driving force for protein/CNT binding [17, 18], the significance of electrostatic interactions in protein binding with single- and multi-walled CNTs has also been reported [19–21]. To evaluate the efficacy of EUF CNT-polymer composite membranes in improving membrane performance in protein processing, a study of the effects of upstream protein/CNT binding on the protein transmission through the downstream UF membrane is required.

In the present work, we investigate the effects of protein-protein, protein-membrane electrostatic interactions on the permeate flux and protein sieving during single and binary protein ultrafiltration/electroultrafiltration using both unmodified polsulfone (PS-35) membranes and electrically conductive PVA-CNT/PS-35 (denoted as PVA-CNT) membranes (30 kDa MWCO) which acted as the cathode. Model proteins of similar size but different charge properties, α-lactalbumin (αLA; molecular weight, MW: 14.2 kDa; isoelectric point, pI = 4.5) and hen egg-white lysozyme (HEL; MW: 14.3 kDa; pI = 11.4), are chosen for the crossflow EUF at pH 7.4 and 4 mM ionic strength. Effects of membrane fouling on the transient flux and protein transmission are studied. Evaluation of the global zeta potential through the membranes are performed before and after protein EUF to probe the interactions of the particle and the membrane surface which contribute to fouling [22, 23]. Visualization of the surface coverage with and without an applied electrical potential is obtained using scanning electron microscopy to investigate protein fouling directly above the conductive thin film. We demonstrate that at moderate applied potentials of −4.6 V (vs. Ag/AgCl), separation of binary protein solutions could be temporarily enhanced although primarily through modified preferential adsorption.

## Materials and methods

### Materials and preparation

Unmodified commercial polysulfone UF membranes (PS-35, Nanostone Water, Inc., Oceanside, CA, USA) with a reported average pore size of 35 nm and molecular weight cutoff of 30 kDa were used for the single and binary protein UF/EUF experiments. PS-35 membranes were also were deposited with a CNT-polymer thin film to form electrically conductive composite membranes for use in the EUF experiments. Multi-walled, carboxyl-functionalized CNTs with a reported outer diameter of 13 – 18 nm, length of 3 – 30 μm, functional group content of 7.0%, and purity of > 99 wt% (SKU 030303, Cheap Tubes, Inc., Cambridgeport, VT, USA) were used to modify the UF membranes. A solution of 50 wt% glutaraldehyde (G151-1, Thermo Fisher Scientific, Inc., Waltham, MA, USA), 146,000 – 186,000 MW poly(vinyl-alcohol) (SKU 363065, Sigma-Aldrich, Corp., St. Louis, MO, USA), and sodium dodecylbenzenesulfonate (SKU 289957, Sigma-Aldrich, Corp., St. Louis, MO, USA) were used for preparation of the CNT suspension.

Experiments were performed with α-lactalbumin whey protein isolate (Agropur, Inc., Eden Prairie, MN, USA) and hen egg-white lysozyme (L6876, Sigma-Aldrich, Corp., St. Louis, MO, USA). Table 1 lists the physicochemical properties of the model proteins used. Protein solutions were prepared by adding a predetermined mass (AG204 DeltaRange, Mettler-Toledo, Inc., Columbus, OH, USA) of lyophilized protein in the appropriate volume of deionized (DI) water. For single protein filtration experiments, the feed solutions contained 0.1 g/L protein concentration; for binary protein filtration experiments, the feed solutions contained 0.1 g/L protein concentration of each protein component. All solutions were prepared using DI water with resistivity of 18.2 MΩ cm at 25 °C (Model 50132370, Barnstead Micropure UV/UF; Thermo Fisher Scientific; Waltham, MA, USA). The ionic strength of the feed solutions was adjusted by adding the appropriate amount of NaCl into the feed solution to achieve 1 mM salt concentration. Besides NaCl, 1 mM of Na_2_HPO_4_ was added as a buffer to yield a final solution ionic strength of 4 mM. The solution pH was adjusted to 7.4 by dropwise additions of 0.1 M HCl or NaOH solution and measured with a pH meter (Model 13-641-253, Thermo Scientific Orion 720A+, Thermo Fisher Scientific, Inc., Waltham, MA, USA). The protein UF experiments were conducted with protein feed solutions at room temperature (25 °C).

**Table 1.** Physicochemical properties of model proteins used in ultrafiltration experiments.

| Characteristics | Hen egg-white lysozyme (HEL) | α-lactalbumin (αLA) |
| --- | --- | --- |
| Molecular weight (kDa) | 14.3 [24] | 14.2 [25] |
| Isoelectric point | 11.0 [26] | 5.2 [27] |
| Net charge at pH 7.4 | 8.0<sup>a</sup> | -7.0<sup>a</sup> |
| Stokes radius (nm) | 2.09 [28] | 2.02 [29] |
| Diffusion coefficient (10<sup>-10</sup> m<sup>2</sup>/s) | 1.18 [30] | 1.06 [31] |
| Hydrophobicity (Cal/residue) | 970 [32] | 1150 [32] |
| Surface hydrophobicity | 1.66 [33] | 7.49 [33] |

<sup>a</sup> Calculated from PDB2PQR 2.0.0/PROPKA 3.0 from PDB file (1DPX, 1HFZ).
<sup>b</sup> Calculated from amino acid sequence.
<sup>c</sup> Determined as the retention coefficients from hydrophobic column chromatography.

### Membrane fabrication

The fabrication process for the PVA-CNT m+embranes followed the method previously reported by Dudchenko et al. [6] with adjustments to the cross-linking procedure. Briefly, PS-35 UF membranes were thoroughly wetted with DI water prior to modification. For preparation of the carbon nanotube suspension, 0.01 wt% CNT-COOH powder and 0.1 wt% dodecylbenzenesulfonic acid were suspended in DI water using a horn sonicator (Branson, Danbury, CT, USA). A 3:1 ratio of 1 wt% PVA to CNT-COOH solution was pressure-deposited onto the PS-35 polysulfone UF membranes. Cross-linking between PVA and CNT-COOH was achieved through immersion in 1 g/L glutaraldehyde and 0.37 g/L of hydrochloric acid at 90 °C for 1 h. The membranes were subsequently removed from the heated solution, dried at 90 °C for 5 min, and then stored at room temperature before use.

### Crossflow electroultrafiltration module

All UF/EUF experiments were conducted in a crossflow system (XX42LSS11, MilliporeSigma, Burlington, MA, USA) with a customized flow cell unit (Fig 2). The experiments were conducted using a crossflow filtration configuration in which the feed solution was siphoned into the retentate tank (500 mL feed volume) from a secondary reservoir via vacuum as the permeate was leaving the system. Hence, the feed volume remained constant. The permeate solution was collected while the retentate was recycled to the primary feed tank.

**Fig 2.**
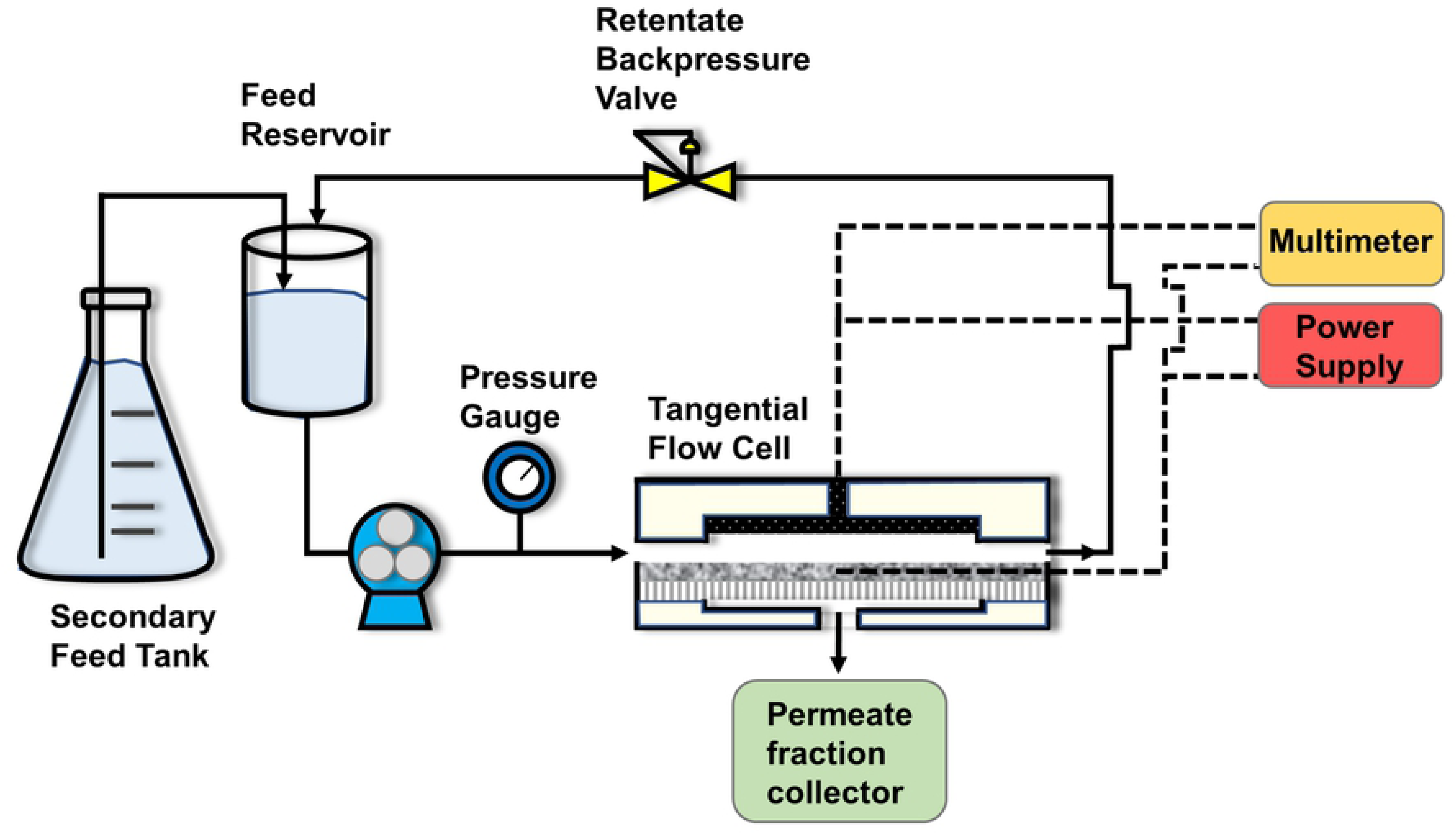
Experimental crossflow EUF system flow diagram. Solid lines represent solution tubing; dashed lines represent electrical wiring.

The custom-built flow cell features a carbon fiber mesh counter-electrode in the top chamber with an attached carbon fiber mesh wire extending out of the top compartment for electrical connectivity. The membrane was stabilized between the two chambers of the flow cell with a thin, porous plastic mesh placed inside each chamber to provide mechanical support and prevent electrical contact with the counter electrode. The top and bottom compartments of the flow cell are each 8.62 cm in width, 5.00 cm in length, and 0.03 cm, in height with an effective membrane surface area of 43.10 cm^2^ and cross-sectional chamber area of 0.26 cm^2^. All EUF experiments were conducted with the membrane functioning as the cathode (negatively charged) and the carbon fiber mesh electrode functioning as the anode. Electrical contacts were made at the carbon fiber mesh wire and the two sides of the UF membrane in proximity to the feed inlet and retentate outlet. An external electric field was applied between the electrically conductive UF membrane and the carbon fiber mesh electrode using a direct current output power supply (LLS-4040, Lambda Electronics, Inc., Melville, NY, USA) under constant voltage operation. The relative potential (vs. Ag/AgCl reference electrode) was determined using a potentiostat (Gamry Instruments, Warminster, PA, USA). The potential difference between the carbon fiber mesh electrode and the PVA-CNT composite membrane was monitored during the beginning and end of the protein EUF experiments using a digital multimeter (77 Series II, Fluke, Corp., Everett, WA, USA).

### Experimental procedure

#### Operating conditions

Prior to the protein EUF experiments, the PVA-CNT composite membranes were compacted at 30 psi using DI water with resistivity of 18.2 MΩ cm (Model 50132370, Barnstead Micropure UV/UF; Thermo Fisher Scientific; Waltham, MA, USA). All filtration experiments were performed with protein feed solutions at 0.1 g/L concentration of each protein component, solution pH of 7.4, and ionic strength of 4 mM. The single protein filtration experiments were performed at 1 psi constant applied transmembrane pressure (*ΔP*); 555 s^-1^ crossflow shear rate; and 9.33 h duration. 0 V and 9 V total cell potentials were applied to the membrane/counter electrode with the membrane functioning as the cathode. The binary protein filtration studies were performed with the same experimental flow conditions with 5.33 h of operation at 0 V, 5 V, and 9 V cell potentials. The relative potentials on the membrane (vs. Ag/AgCl reference) at 0 V, 5 V, and 9 V total cell potentials were determined to be 0 V, −2.5 V, and −4.6 V, respectively.

The membrane hydraulic permeability was evaluated without an applied electric field, *E_y_*, before and after the protein filtration experiments:

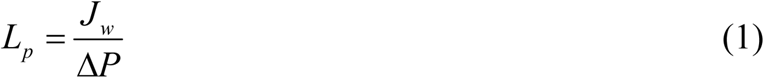

where *J_w_* is the volumetric water flux (volumetric flow rate per membrane area) and *ΔP* is the transmembrane pressure driving force. The permeate solution was collected into test tubes using a fraction collector (Retriever 500 and Retriever II; Teledyne Technologies, Inc.; Thousand Oaks, CA, USA), and the permeate flux (*J_v_*) was recorded. After the experiments, the membranes were removed from the system, rinsed with DI water, immersed in 0.4 wt% Terg-a-zyme enzymatic detergent solution at 40 °C for 1 h, rinsed thoroughly with DI water, and stored in DI water for at least 8 h before reuse. The cleaning procedure followed an optimal enzymatic cleaning procedure for polysulfone membranes [34]. The crossflow filtration system was also cleaned with the enzymatic detergent followed by a rinse with DI water. All crossflow filtration experiments were completed in at least duplicate. The protocol for evaluating the protein concentrations is provided in the Supporting Information A.4.1. The observed sieving coefficient was calculated:

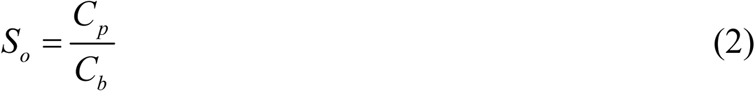

where *C_p_* is the permeate protein concentration and *C_b_* is the bulk protein feed concentration. From the permeate flux and permeate protein concentration, the mass flux (*N_p_*) was also calculated:

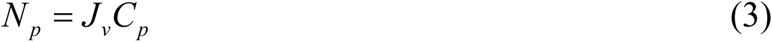

#### Characterization of membranes

PVA-CNT membrane surfaces were imaged using scanning electron microscopy (SEM; Nova NanoSEM 450, Thermo Fisher Scientific; Waltham, MA, USA) to assess fouling on top of the composite membranes. Membrane samples were affixed to carbon conductive tapes on SEM stubs and sputter coated with a Pt/Pd target for 60 s (Sputter coater 108 Auto, Cressington Scientific Instruments Ltd., Watford, England, UK) prior to imaging with SEM. An estimation of the average pore size of the conductive network layer from the SEM images of the membrane surface was performed using the image processing software, ImageJ v1.52e [35].

Streaming potential across the membrane pore walls (*Ψ_p_*) was determined at various pressures using a custom device (Supporting Information A.4.2) [36]. The streaming potential device was filled with 10 mM NaCl solution at pH 7.4 with Ag/AgCl electrodes (Sigma-Aldrich, Corp., St. Louis, MO, USA) attached to each side of the membrane and symmetrically aligned with one another. The changes in streaming potential upon alterations in the hydraulic pressure were detected using a voltmeter (77 Series II Multimeter, Fluke, Corp., Everett, WA, USA).

A three-layer porous structure is considered where layer 1 represents the support layer of the PS-35 membrane, layer 2 represents the skin layer of the PS-35 membrane, and layer 3 represents the deposited PVA-CNT layer. To estimate the relative contributions of each layer (*k*) to the global zeta potential through the membrane, the pore radius, *a^(k)^*, and length, *l^(k)^*, are considered. The length fraction of the support layer, *X*, is defined as

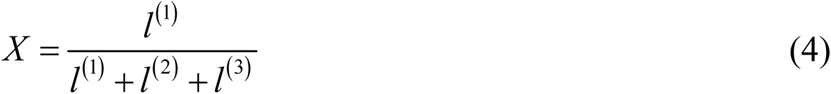

#### Statistical analysis

Two-way analysis of variance (ANOVA) followed by *post hoc* Tukey’s multiple comparisons tests were performed using Graphpad Prism version 6.01 (Graphpad Software, La Jolla, CA, USA) to assess the statistical differences among the weighted factor means at a significance level of *p* < 0.05.

## Results and discussion

### PVA-CNT membrane characterization

From the cross-sectional SEM image of the PVA-CNT/PS-35 composite membrane (S3 Fig), the thickness of the PVA-CNT thin film was determined to be approximately 6 μm compared to the 165 µm thickness of the commercial polysulfone membrane which agrees with previously reported measurements [5, 11]. With an approximate 10 µm thickness of the selective layer [5], the length fraction of the support layer, *X*, was determined to be 0.91. Based on image analysis of SEM images of the PVA-CNT surface using the NIH ImageJ software (minimum size of 0.007 µm^2^, circularity parameter of 0.01-1), the typical pore size of the PVA-CNT network (assuming circular pores) was determined to be 475 nm (S5 Fig). The results indicate that the pore size of the conductive network is significantly larger than that of the underlying selective layer of the polysulfone membrane.

### Membrane cleaning

The PVA-CNT membranes exhibit variability in the initial hydraulic permeability, ranging from approximately 8.6 – 14.5 × 10^-10^ m s^-1^ Pa^-1^ which may be due to both the commercial manufacturing process for the PS-35 polysulfone membranes as well as the procedure for deposition of the PVA-CNT layer. Six PS-35 membranes were randomized and used for the single and binary protein UF studies. Two PVA-CNT membranes were used for the single protein EUF studies: one for the HEL studies and the other for the αLA studies. Another two PVA-CNT membranes were used for the binary protein EUF studies: one for each set of trials (0 V, −2.5 V, - 4.6 V applied potentials to the membrane vs. Ag/AgCl).

The initial hydraulic permeability of uncoated PS-35 membranes were fully restored following each filtration experiment through chemical cleaning which involved soaking the membranes in solution containing the Terg-a-zyme enzymatic cleaning agent. While the chemical cleaning process was also effective for fully restoring PVA-CNT membrane performance following single protein EUF of αLA, the procedure was only able to restore approximately 70 – 95% of the initial hydraulic permeability of the corresponding virgin PVA-CNT membranes following each run for single protein EUF of HEL and binary protein EUF of αLA and HEL. Although the PVA-CNT membrane that was reused following chemical cleaning for single protein EUF of αLA exhibited no loss in hydraulic permeability after four cycles of filtration/cleaning, the PVA-CNT membrane for single protein EUF of HEL had approximately 80% restoration efficiency after three cycles and 60% restoration efficiency after six cycles which indicated irreversible fouling. For binary protein EUF, the average restoration efficiency following three cycles was approximately 85%. The observed restoration efficiencies of the PVA-CNT membrane suggest hydrophobic and electrostatic interactions between the charged proteins and the hydrophobic, negatively charged CNT network of the deposited porous film play critical roles in contributing to irreversible membrane fouling. Although α-lactalbumin is a homologue of lysozyme, the two model proteins have different charge properties; lysozyme is positively charged at the experimental pH of 7.4, whereas α-lactalbumin is negatively charged. Moreover, the surface hydrophobicity of HEL is greater than that of αLA (Table 1). Therefore, it is expected that protein adsorption following EUF of HEL is greater than that following EUF of αLA due to the enhanced hydrophobic and electrostatic interactions leading to irreversible fouling. However, the effect of the applied electrical potentials during electroultrafiltration on the efficiency of PVA-CNT membrane restoration is unclear; a detailed investigation into the impact of applied potentials on the CNT-deposited composite ultrafiltration membranes is beyond the scope of the current study.

### Permeate flux

During electroultrafiltration, the application of an electric field leads to Joule heating resulting in the rise of the feed temperature. In addition, application of a high electric potential to the electrically conducting CNT-polymer composite membrane surface, with the membrane serving as the cathode, results in water electrolysis and hydroxide ion formation along the membrane surface. Therefore, in order to reduce Joule heating and electrolysis effects, the maximum applied constant DC potential was restricted to 4.5 V (vs. Ag/AgCl) which would amount to a negligible change in temperature of the feed (< 2 °C) [37–39].

Due to the different initial fluxes of the membranes, the normalized permeate flux (*J_v_ / J_0_*), the ratio of permeate flux during the filtration process to the initial permeate flux at the beginning of the filtration, was evaluated. Since the transmembrane pressure was held constant, a decrease in the permeate flux corresponds to membrane fouling. Fig 3A-C show the normalized permeate flux during single protein and binary protein studies for constant pressure (1 psi) UF/EUF of lysozyme and α-lactalbumin (0.1 g/L of each protein species, 4 mM ionic strength, pH 7.4) using unmodified PS-35 membranes and electrically conductive PVA-CNT ultrafiltration membranes. For UF of single protein HEL, the steady state normalized permeate fluxes for the PS-35 and PVA-CNT composite membranes following 9.33 h of filtration are 0.66 ± 0.03 and 0.45 ± 0.04, respectively (Fig 3A). Without an applied electric potential, the PVA-CNT layer still carried negative charges at pH 7.4 due to the carboxyl groups at the termini of the multi-walled CNT [21]. The reduction in permeate flux with the PVA-CNT composite membrane compared to the PS-35 (without the PVA-CNT layer) may be due to fouling from the electrostatic attractions of the net positively charged HEL (pI: 11.0) and the negatively charged PVA-CNT layer at the experimental pH of 7.4. Conversely, the presence of the PVA-CNT layer resulted in an increase in the steady state normalized permeate flux for single protein UF of αLA from 0.61 ± 0.08 to 0.85 ± 0.03 due to the electrostatic repulsion between the net negatively charged αLA (pI: 5.2) and the negatively charged PVA-CNT layer (Fig 3B).

**Fig 3.**
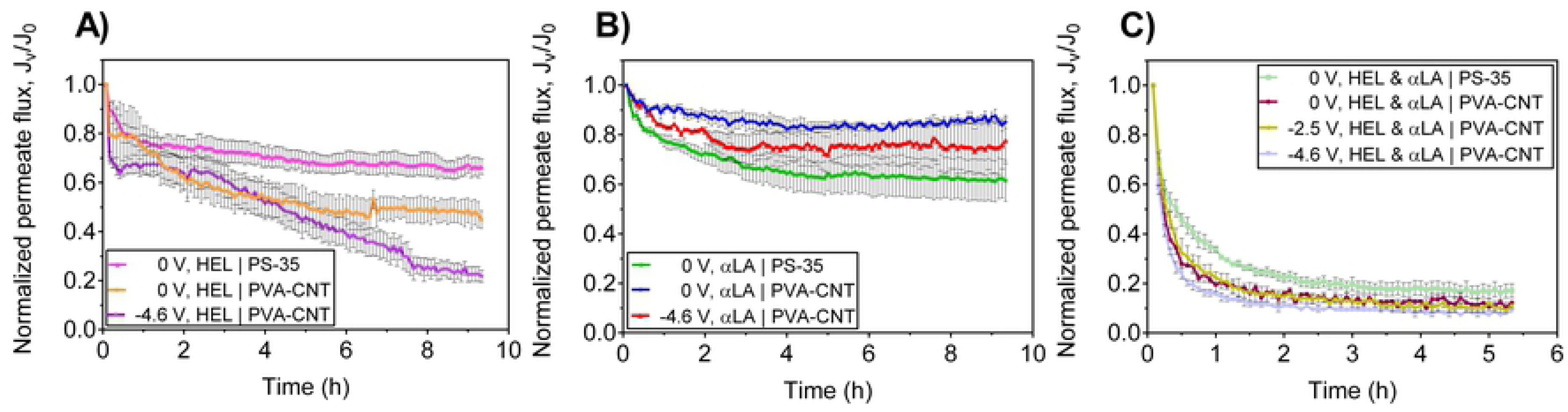
Carbon nanotube-polymer thin film deposition and electric potential significantly affect the steady state permeate flux and rates of permeate flux decline. Normalized permeate flux during UF/EUF of single protein and binary protein solutions at TMP of 1 psi with different cathodic potentials (vs. Ag/AgCl) applied to the PS-35 and PVA-CNT membranes: A) single protein solution of 0.1 g/L HEL (n = 3), B) single protein solution of 0.1 g/L αLA (PS-35, n = 3; PVA-CNT, n = 2), and C) binary protein solution containing 0.1 g/L HEL and 0.1 g/L αLA (PS-35, n = 3; PVA-CNT, n = 2). Error bars represent the standard error of the weighted mean. Electrostatic interactions between the proteins and the carboxylated multiwalled CNTs significantly affect the normalized steady state permeate flux during single protein UF and the rate of flux decline during binary protein UF. The abrupt changes in permeate flux during single protein EUF suggest the application of an external electric potential influences the extent of protein adsorption on the PVA-CNT layer.

Single protein studies with the cathodic PVA-CNT membrane exhibited flux behavior not typical of flux decline in UF processes (Fig 3A-B). For both cases of crossflow EUF of single protein solutions, the normalized permeate flux at −4.6 V vs. Ag/AgCl decreased abruptly following around 2 to 3 h. The atypical flux decline during EUF with an applied electric field suggests high protein adsorption at the PVA-CNT membrane surface and development of a filter cake. Significant protein adsorption and permeate flux decline is expected for EUF of HEL due to the electrostatic attraction between the negatively charged, carboxylated, multiwalled CNTs and the net positively charged HEL protein. However, the net negatively charged α-lactalbumin contains positively charged regions (S7 Fig) which may also bind to the CNTs, resulting in the decline of the observed permeate flux during EUF of αLA. The steady state normalized permeate flux for single protein EUF of HEL and single protein EUF of αLA are 0.22 ± 0.02 and 0.78 ± 0.09, respectively, corresponding to a 51% and 8% reduction with an applied cell potential of −4.6 vs. Ag/AgCl compared to values for UF of HEL and αLA without an applied cell potential (Fig 3A-B). The decrease in the steady state normalized permeate flux is more significant for single protein filtration of the net positively charged lysozyme than for the net negatively charged α-lactalbumin at pH 7.4 due to the respective attractive and repulsive electrostatic interactions with the negatively charged PVA-CNT membrane layer. The more pronounced reduction in permeate flux for single protein HEL compared to single protein αLA may also be a result of pore narrowing and plugging due to adsorption of HEL on carboxylated multiwalled CNTs [19].

A significant reduction in permeate flux was observed in the binary protein studies using the PS-35 membrane, and the PVA-CNT composite membrane at 0 V, −2.5 V, and −4.6 V vs. Ag/AgCl, with steady state normalized permeate flux values of 0.17 ± 0.03, 0.12 ± 0.01, 0.11 ± 0.01, and 0.09 ± 0.01, respectively (Fig 3C). For EUF of the binary protein solution of αLA and HEL, the PVA-CNT composite membrane resulted in an increased rate of flux decline compared to the PS-35 membrane which may be attributed to binding of HEL and αLA-HEL aggregates to the PVA-CNT layer due to electrostatic interactions. The application of the electric potential did not influence the long-term flux performance during binary protein EUF.

### Protein sieving and mass flux

Fig 4 shows the transient response in observed sieving and mass flux during UF/EUF of single protein feed solutions of 0.1 g/L lysozyme (Fig 4A and Fig **4C**) and 0.1 g/L α-lactalbumin (Fig 4B and Fig **4**D) with and without a constant cathodic potential of −4.6 V vs. Ag/AgCl applied to the PVA-CNT/PS-35 membranes. In the incipient stage of single protein HEL with and without the applied potential, the PS-35 and the PVA-CNT membrane exhibit near complete rejection of lysozyme (Fig 4A and Fig **4**C). The low HEL transmission through the PS-35 membranes may be attributed to the initial adsorption of the lysozyme solutes on the hydrophobic, negatively charged surface of the polysulfone membrane. The initial stage of lysozyme adsorption is further increased with the additional deposited conductive layer on the PVA-CNT membranes. However, single protein UF of αLA using the PS-35 membrane exhibited higher observed sieving and mass flux at the beginning of operation likely due to the reduced initial adsorption on the membrane as a result of the electrostatic repulsion of the net negatively charged αLA and the unmodified polysulfone membrane which also possesses a negative charge at the operating pH (Fig 4B and Fig **4**D) [22, 40]. During single protein UF of αLA without an applied potential using the PVA-CNT membrane, near complete rejection of αLA was observed during the beginning of operation. This may be a result of the initial adsorption of the αLA solutes on the CNT network. The adsorption of αLA on the CNT network during single protein EUF, however, is mitigated with an applied potential due to the electrostatic repulsion of the net negatively charged αLA and the negatively charged PVA-CNT composite membrane.

**Fig 4.**
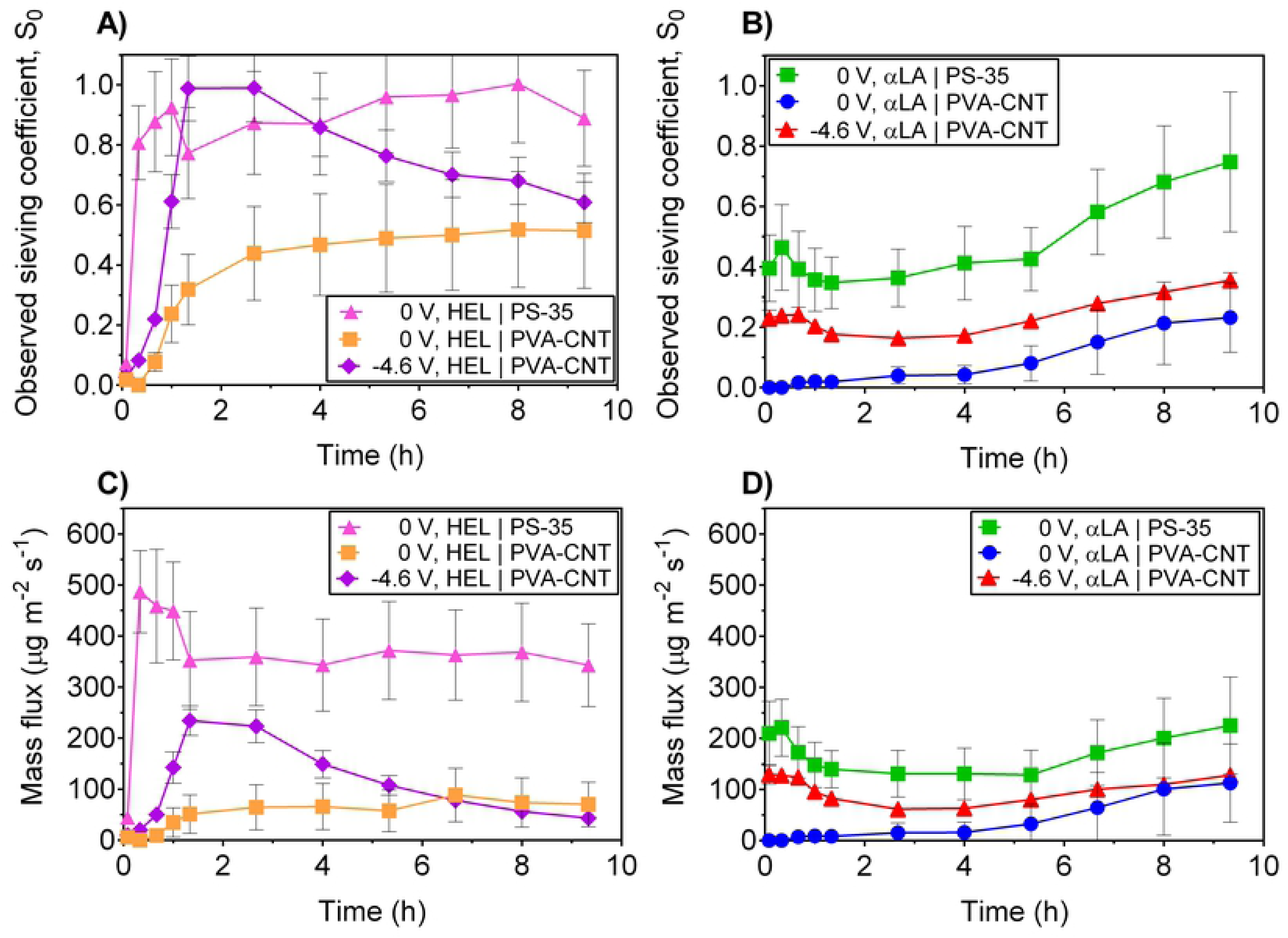
Temporary enhancement in protein sieving with an applied potential is seen during electroultrafiltration of single protein solutions. Observed sieving coefficient and mass flux during UF/EUF of single protein solutions at TMP of 1 psi at different cathodic potentials (vs. Ag/AgCl) applied to the PS-35 and PVA-CNT membranes: A) and C) single protein solution of 0.1 g/L HEL (n = 3), B) and D) single protein solution of 0.1 g/L αLA (PS-35, n = 3; PVA-CNT, n = 2). Error bars represent the standard error of the weighted mean. Application of an external electric potential results in temporary improvement in sieving and mass flux during both single protein of HEL and αLA.

For the single protein UF experiments using the PS-35 membrane without the PVA-CNT layer, the observed sieving coefficient at the end of the 9.33 h filtration run was 0.9 ± 0.16 for UF of HEL and 0.7 ± 0.23 for UF of αLA (Fig 4A-B). Correspondingly, the mass fluxes at the end of the single UF experiments for HEL and αLA were 340 ± 81 µg m^-2^ s^-1^ and 220 ± 95 µg m^-2^ s^-1^, respectively (Fig 4C-D). The apparent rejections of the 14 kDa proteins were considerably high for a nominal 30 kDa cutoff membrane which may be due to a combination of electrostatic forces rejecting the proteins from traversing the membrane pores, and pore narrowing/blockage from protein fouling with the hydrophobic PS-35 membrane. At the operational pH of 7.4, the HEL protein is electropositive while the αLA protein and the polysulfone membrane are electronegative [24]. Consequently, the observed sieving of αLA is lesser than that of HEL during single protein UF following 9.33 of filtration due to the electrostatic repulsive force acting between the αLA proteins and the PS-35 membrane. With the addition of the PVA-CNT layer on the membrane, the observed sieving for HEL following 9.33 h of filtration decreased 44% to 0.5 ± 0.19 while the observed sieving for αLA decreased 71% to 0.2 ± 0.11 (Fig 4A-B). The corresponding mass fluxes at the end of the single protein UF experiments were 70 ± 44 µg m^-2^ s^-1^ for HEL filtration and 110 ± 76 µg m^-2^ s^-1^ for αLA filtration (Fig 4C-D). The reduction in observed sieving and mass flux with the deposited CNT layer even without an applied electric potential is likely a result of the high protein affinity of the carboxylated multiwalled CNTs [19, 21].

During EUF of single protein lysozyme at an applied potential of −4.6 V vs. Ag/AgCl, the observed sieving coefficient increased from the initial value of 0.03 ± 0.011 to peak values of 1.0 ± 0.11 following filtration for 1.33 h and 2.67 h (Fig 4A). Correspondingly, the mass flux increased from 10 ± 3.6 µg m^-2^ s^-1^ at the initial timepoint following 5 min of EUF to peak values of 230 ± 29 µg m^-2^ s^-1^ at 1.33 h and 220 ± 32 µg m^-2^ s^-1^ at 2.67 h (Fig 4C). Therefore, single protein EUF of HEL using the PVA-CNT composite membrane with an application of −4.6 V cell potential vs. Ag/AgCl yielded an approximate 3-fold enhancement in the observed sieving and 4-fold in the mass flux over single protein UF of HEL without an applied potential. Application of a potential during single protein EUF of HEL leads to electrophoretic migration and electrostatic attraction of HEL proteins toward the membrane. Hence, the HEL concentration at the membrane surface increases and more HEL proteins are transported through the membrane resulting in temporary increases in both the observed sieving and mass flux. Interestingly, for single protein EUF of the similarly sized but net negatively charged α-lactalbumin using the PVA-CNT membrane, application of an applied potential of −4.6 V vs. Ag/AgCl also yielded a marked enhancement in both the observed sieving and mass flux compared to runs with no applied potential (Fig 4B and Fig **4**D). The PVA-CNT membrane without an applied potential exhibited near total rejection of αLA during the first 2.67 h of EUF. Use of the PVA-CNT membrane with application of an electric potential (−4.6 V vs. Ag/AgCl), however, yielded an approximate 15-fold improvement in the observed sieving coefficient and mass flux with values of 0.24 ± 0.013 and 120 ± 10.5 µg m^-2^ s^-1^ at 0.66 h, respectively. Although αLA has a net negative charge, calculations of the electrostatic potential and examination of the protein charge distribution reveal positively charged regions on the protein surface (S7 Fig). Adsorption of αLA on the PVA-CNT membrane surface occurs with the αLA proteins oriented such that the complementary, positively charged area faces the adsorbing PVA-CNT substrate [41]. In this way, non-adsorbed αLA proteins with a net negative charge at pH 7.4 may experience electrostatic repulsive forces from the exposed negatively charged protein layer adsorbed on the membrane surface which lead to higher observed sieving and mass flux at the early stage of electroultrafiltration. However, it is important to emphasize that the improvements in observed protein sieving and mass flux are temporary due to continued protein adsorption on the membrane surface.

As shown in Fig 4, although application of the electric potential during single protein EUF using the PVA-CNT membrane led to higher observed sieving and mass flux during the initial stage of filtration, the improvement was temporary. For single protein EUF of HEL at an applied potential of −4.6 V vs. Ag/AgCl, the observed sieving coefficient and mass flux began decreasing from their peak values following 2.67 h of EUF to values of 0.61 ± 0.068 and 43 ± 5.7 µg m^-2^ s^-1^, respectively, after 9.33 h (Fig 4A and Fig **4**C). Application of an electric potential during single protein EUF of αLA also yielded an improvement in the observed sieving and mass flux following 2.67 h of EUF but exhibited convergence towards observed sieving coefficient and mass flux of 0.35 ± 0.025 and 128 ± 3.6 µg m^-2^ s^-1^, respectively, following 9.33 h of EUF (Fig 4B and Fig **4**D). The timepoints of the inflections for observed protein sieving and mass flux during EUF of the two single protein solutions in Fig 4 correspond well with the respective times at which the sharp decline in normalized permeate fluxes were observed in Fig 3A-B. The dramatic decline in permeate flux for both cases of single protein EUF suggest significant pore plugging from the retentate cake formation led to the pronounced reductions in protein transmission through the PVA-CNT membrane. Consequently, at extended operation times, single protein EUF using the PVA-CNT membrane with an application of −4.6 V vs. Ag/AgCl does not yield a significant improvement in observed sieving and mass flux over UF without an applied potential.

Binary protein UF/EUF of feed solutions containing αLA and HEL (0.1 g/L of each species) were also performed using the PS-35 and PVA-CNT membranes (Fig 5). Comparisons of the transient changes in observed sieving and mass flux between the unmodified PS-35 membrane and the PVA-CNT composite membrane are shown in Fig 5A and Fig 5C. The effects of the applied potentials (0 V, −2.5 V, and −4.6 V vs. Ag/AgCl) on the observed sieving and mass flux of each protein species during binary protein EUF using the PVA-CNT membrane are shown in Fig 5B and Fig 5D. The unmodified PS-35 membranes were not highly selective in separating αLA and HEL during ultrafiltration of the binary protein solution (Fig 5A and Fig 5C). However, use of the PVA-CNT composite membrane consisting of the additional conductive, porous layer even without an applied potential resulted in nearly full rejection of the positively charged lysozyme during the entirety of the EUF experiments and displayed the highest protein selectivity at 20 min. For the binary protein EUF studies, nearly full rejection of the positively charged HEL was also achieved at cathodic potentials of −2.5 V and −4.6 V vs. Ag/AgCl. Compared to the observed sieving coefficient when no potential is applied, the observed sieving coefficient of the negatively charged αLA, however, was enhanced by approximately 36% and 69% at the 20 min period of operation with application of potentials of −2.5 V and −4.6 V, respectively (Fig 5B). Correspondingly, the mass flux of αLA at 20 min of binary protein EUF was enhanced by approximately 127% and 105% at applied potentials of −2.5 V and −4.6 V vs. Ag/AgCl, respectively (Fig 5D). As was observed in the single protein EUF studies, binary protein EUF with an applied potential resulted in only a temporary improvement in observed sieving and mass flux over ultrafiltration without an applied potential. The reduced duration of effectiveness during binary protein EUF over single protein EUF may be explained by the higher protein concentration of the binary protein feed solution and protein aggregation due to the presence of positively charged HEL and negatively charged αLA at low ionic strength which may contribute to increased fouling.

**Fig 5.**
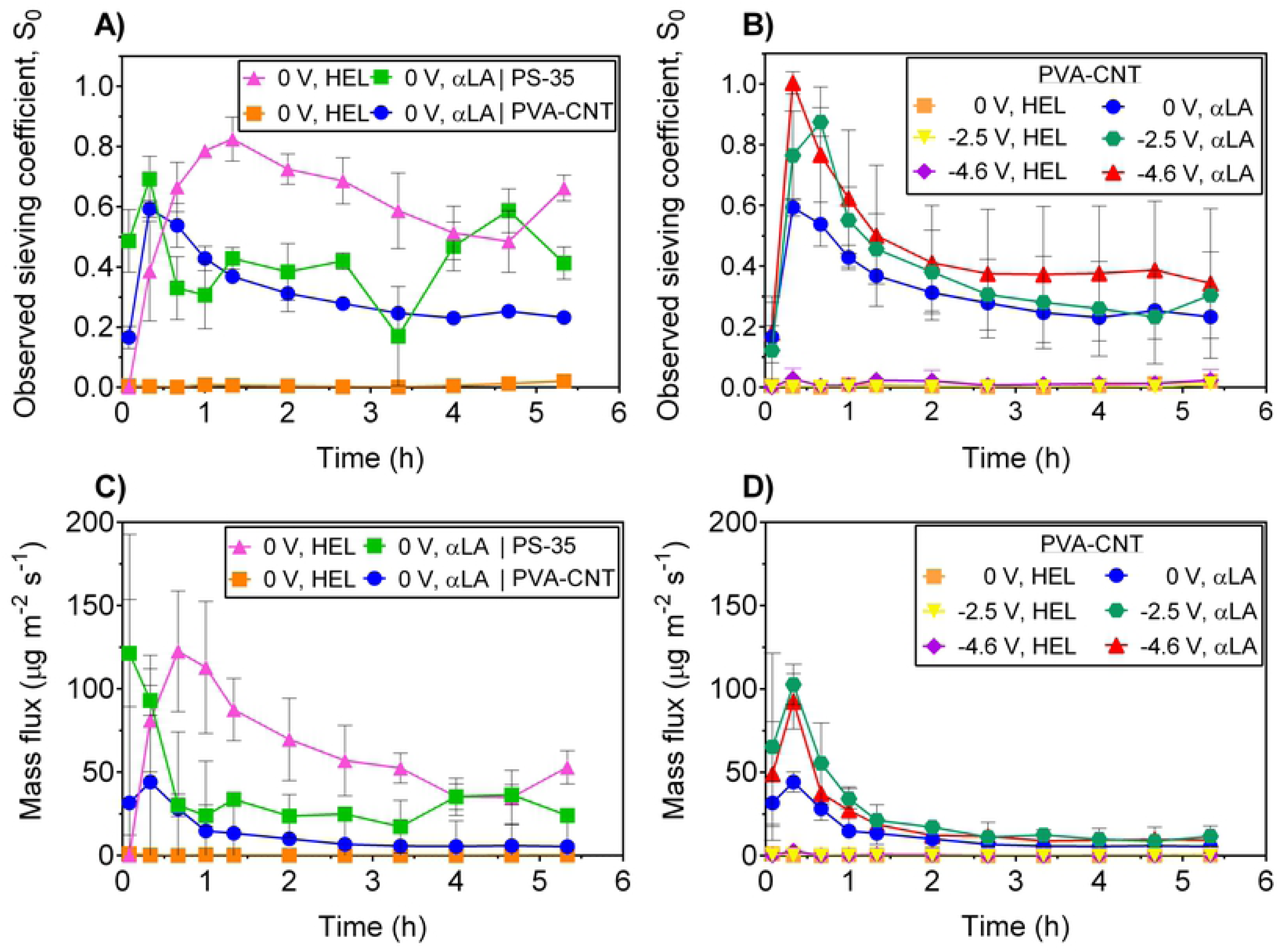
Temporary enhancement in selectivity is seen during initial stage of binary protein electroultrafiltration. Observed sieving coefficient and mass flux during UF/EUF of binary protein solutions at TMP of 1 psi at different cathodic potentials (vs. Ag/AgCl) applied to the PS-35 and PVA-CNT membranes (PS-35, n = 3; PVA-CNT, n = 2): A) and C) Comparison of membranes with (PVA-CNT) and without (PS-35; control) the deposited conductive thin film. B) and D) Comparison of cathodic potentials (vs. Ag/AgCl). Deposition of the PVA-CNT layer on the PS-35 membrane results in temporary separation of binary protein solutions containing species of differing net charges with further enhancement in selectivity upon application of an external electric potential.

A binary protein EUF study was also performed at an applied potential of 2.5 V vs. Ag/AgCl with the PVA-CNT membrane functioning as an anode (S8 Fig) which yielded a reduction in the protein selectivity (*S_o,αLA_/S_o,HEL_*). Although the sieving results from the application of positive, anodic potentials show promise in the tunability of protein selectivity using the PVA-CNT membrane, several studies have indicated CNT-based membranes are unstable under elevated anodic potentials in aqueous environments due to oxidation and degradation of the carbon nanotubes when exposed to hydroxyl radicals generated on the CNT surface [42, 43].

The transient results for the observed sieving and mass flux during single and binary protein EUF highlight the complexity of the transport phenomena for protein electroultrafiltration using an electrically conductive composite ultrafiltration membrane. Transmission of the charged proteins through the CNT-network and the base ultrafiltration membrane is dependent on the contributions from the convective, diffusive, electrophoretic, and electrostatic forces. Application of an electric potential during both single protein EUF of αLA and single protein EUF of HEL as well as for binary protein EUF of αLA and HEL using the PVA-CNT membrane yielded temporary enhancements in the observed sieving and mass flux. However, as EUF proceeded, the development of the filter cake of retained protein resulted in pore narrowing and plugging which contributed to the reduction in observed sieving. Both the net positively charged HEL and net negatively charged αLA proteins exhibited this behavior which is consistent with previous reports of an increase in adsorption of both complementary and oppositely charged proteins to a charged electrode surface upon application of an electric field [41]. The non-ideality in the sieving behavior during EUF and the reversion to the steady state values for the sieving coefficient coincide with the kinetics of overshooting protein adsorption on charged surfaces in previous studies [44–46]. Leading up to the peak in observed sieving, the protein adsorption proceeds at low surface densities with negligible lateral interaction between bound proteins at the membrane surface. The times of the peak in observed sieving for single protein EUF and binary protein EUF were on the order of 0.5 – 2 h and 20 min, respectively, which are consistent with previous reports of the periods of time leading up to the overshoot in adsorbed proteins [44]. The overshoot is typically explained by a transition in the adsorption of proteins from an initial state at low surface densities to a final irreversible state at high surface densities through conformational reorientation of the adsorbed proteins [47, 48]. Protein transmission at the extended stages of EUF is thus highly dependent on the electrostatic interactions between the proteins in the bulk solution and the fouled membrane comprised of adsorbed proteins in their final irreversible state and deposited proteins.

### Membrane protein fouling

Pre- and post-experimental hydraulic permeability data of the conditioned membranes were measured to assess the extent of fouling following the UF/EUF experiments. Single protein EUF of HEL and single protein EUF of αLA using the unmodified polysulfone PS-35 membrane yielded a respective reduction in hydraulic permeability of 46 ± 3% and 31 ± 3% following the experiments. As was observed from the measured permeate flux (Fig 3), the addition of the PVA-CNT thin film on the membrane resulted in greater fouling for single protein EUF of HEL when compared to that for single protein EUF of αLA which may be attributed to the electrostatic and hydrophobic interactions of the protein and the PVA-CNT membrane. At no applied electric field, the reduction in *L_p_* following single protein EUF is more significant for UF of the lysozyme protein solution than for UF of the α-lactalbumin protein solution with observed reductions in *L_p_* of 75.7 ± 0.9% and 25 ± 2%, respectively. A moderate increase in fouling was observed when applying a potential of −4.6 V vs. Ag/AgCl, with a reduction of 80 ± 1% in *L_p_* for the EUF of the single protein lysozyme solution and a decrease of 29 ± 2% in *L_p_* for the EUF of the single protein α-lactalbumin solution. The decrease in hydraulic permeability provides further evidence that membrane fouling was a significant factor in the transient flux and sieving behavior during protein UF with and without an applied electric field. The greater overall reduction in hydraulic permeability for EUF of the HEL solution compared to that for EUF of the αLA solution despite the similar sizes of the protein species indicate the electrostatic attractive forces between the net positively charged HEL and the net negatively charged PVA-CNT membrane result in more significant fouling. For UF/EUF of binary protein solution of 0.1 g/L HEL and 0.1 g/L αLA, the differences in the change in hydraulic permeability among the experiments with the PS-35 membrane at 0 V (vs. Ag/AgCl), PVA-CNT membrane at 0 V, PVA-CNT at −2.5 V, and PVA-CNT at −4.6 V – with respective percent changes in *L_p_* of −90 ± 3%, −95 ± 7%, −95 ± 5%, and −89 ± 1% – were insignificant. Compared to single protein UF/EUF, all cases of binary protein UF/EUF exhibited higher membrane fouling which may be attributed to the higher bulk protein concentration and aggregation of αLA-HEL complexes.

Evaluation of the zeta potential of the PS-35 and PVA-CNT membranes in their virgin and fouled states after protein EUF (at 0 V and 4.6 V potentials vs. Ag/AgCl, 555 s^-1^ crossflow shear rate, 1 psi transmembrane pressure, solution pH 7.4, and 0.1 g/L protein concentration for each species) is provided in Table 2. It is important to note that the apparent zeta potential from the current study represents the global zeta potential through the multiple layers of the composite membrane. The PVA-CNT composite membranes consist of three layers: a thin PVA-CNT layer that provides electrical conductivity, a thin skin layer that provides the membrane with its selectivity, and a much thicker and more porous support layer that provides necessary mechanical strength to the membrane. In such multilayer membranes, the relative contribution of each of the layers to the global streaming potential is dependent on both the physical properties (porosity, pore radius, layer thickness) and electrical properties (zeta potential) of the layers [49, 50]. Syzmczyk et al. coupled streaming potential and permeate flux measurements to characterize the electrokinetic behavior of each layer of alumina membranes and the respective contributions of each of the layers to the global streaming potential [49]. Although a similar methodology is ideal for investigating the charge properties and fouling of the PVA-CNT composite membranes, the individual contributions of each membrane layer could not be elucidated since the thin PVA-CNT layer and the selective layer of the PS-35 membrane were unable to be autosupported. However, under careful interpretation, the measurement of the global zeta potential of the PVA-CNT composite membranes provides critical insights into the extent of membrane fouling during protein EUF.

**Table 2.**
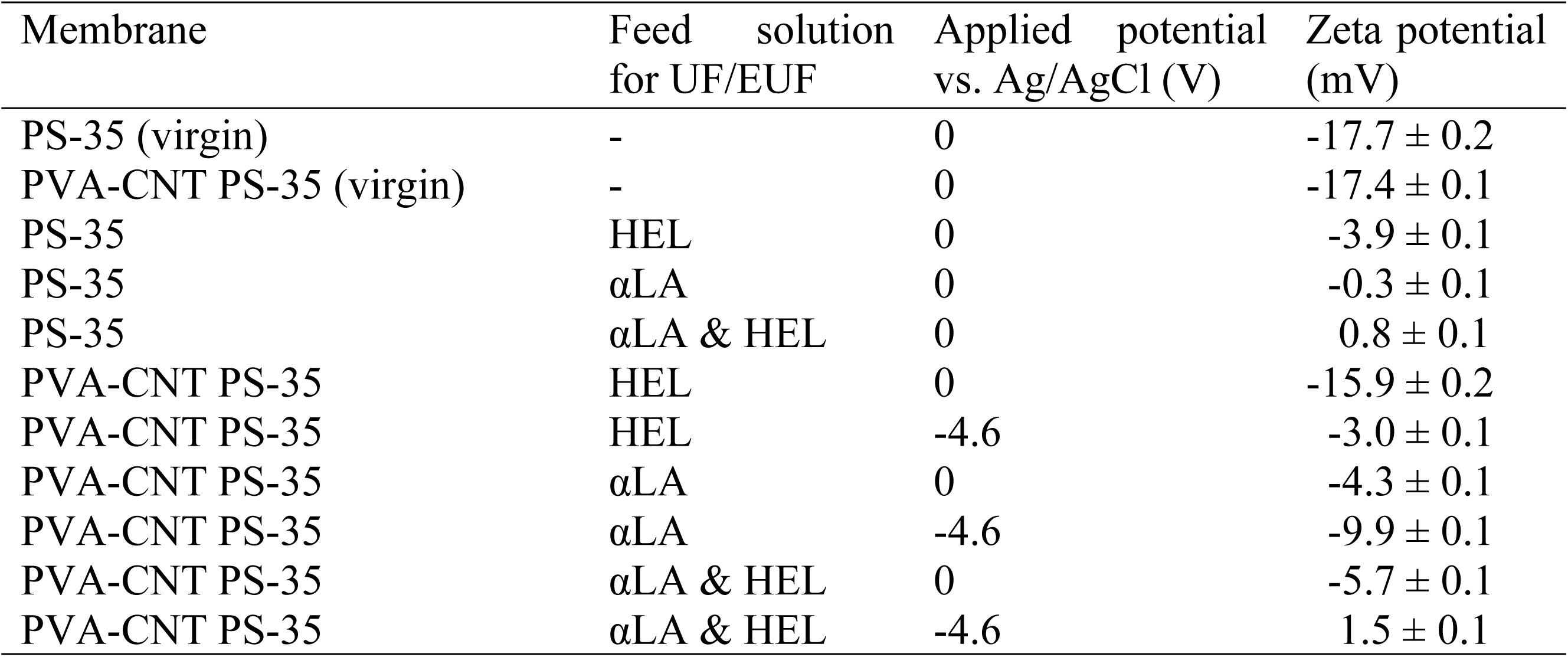
Zeta potential measurements of ultrafiltration membranes after protein ultrafiltration and electroultrafiltration at pH 7.4 and 4 mM ionic strength with the membrane functioning as the cathode.

| Membrane | Feed solution for UF/EUF | Applied potential vs. Ag/AgCl (V) | Zeta potential (mV) |
| --- | --- | --- | --- |
| PS-35 (virgin) | - | 0 | $-17.7 \pm 0.2$ |
| PVA-CNT PS-35 (virgin) | - | 0 | $-17.4 \pm 0.1$ |
| PS-35 | HEL | 0 | $-3.9 \pm 0.1$ |
| PS-35 | αLA | 0 | $-0.3 \pm 0.1$ |
| PS-35 | αLA & HEL | 0 | $0.8 \pm 0.1$ |
| PVA-CNT PS-35 | HEL | 0 | $-15.9 \pm 0.2$ |
| PVA-CNT PS-35 | HEL | -4.6 | $-3.0 \pm 0.1$ |
| PVA-CNT PS-35 | αLA | 0 | $-4.3 \pm 0.1$ |
| PVA-CNT PS-35 | αLA | -4.6 | $-9.9 \pm 0.1$ |
| PVA-CNT PS-35 | αLA & HEL | 0 | $-5.7 \pm 0.1$ |
| PVA-CNT PS-35 | αLA & HEL | -4.6 | $1.5 \pm 0.1$ |

As expected for the hydrophobic polysulfone membrane (PS-35) with a reported isoelectric point between 3.0 – 4.0 [22, 40], the sign of its measured zeta potential in the virgin state was negative (−17.7 ± 0.2 mV) at the experimental pH 7.4 with close agreement to another assessment of the zeta potential of a similar polysulfone membrane [40]. The measured zeta potential of the virgin UF membrane following modification from the deposition of the conductive PVA-CNT layer remained relatively unchanged (−17.4 ± 0.1 mV). At small pore radius ratios (*a^(1)^*/*a^(3)^*) and large length fractions of the support layer (*X*), the thin PVA-CNT layer is therefore expected to have a minimal effect on the electrokinetic behavior of the composite membrane due to the dominant contribution of the pressure drop across the thick PS-35 membrane to the global streaming potential (Fig 6A) [49, 51].

**Fig 6.**
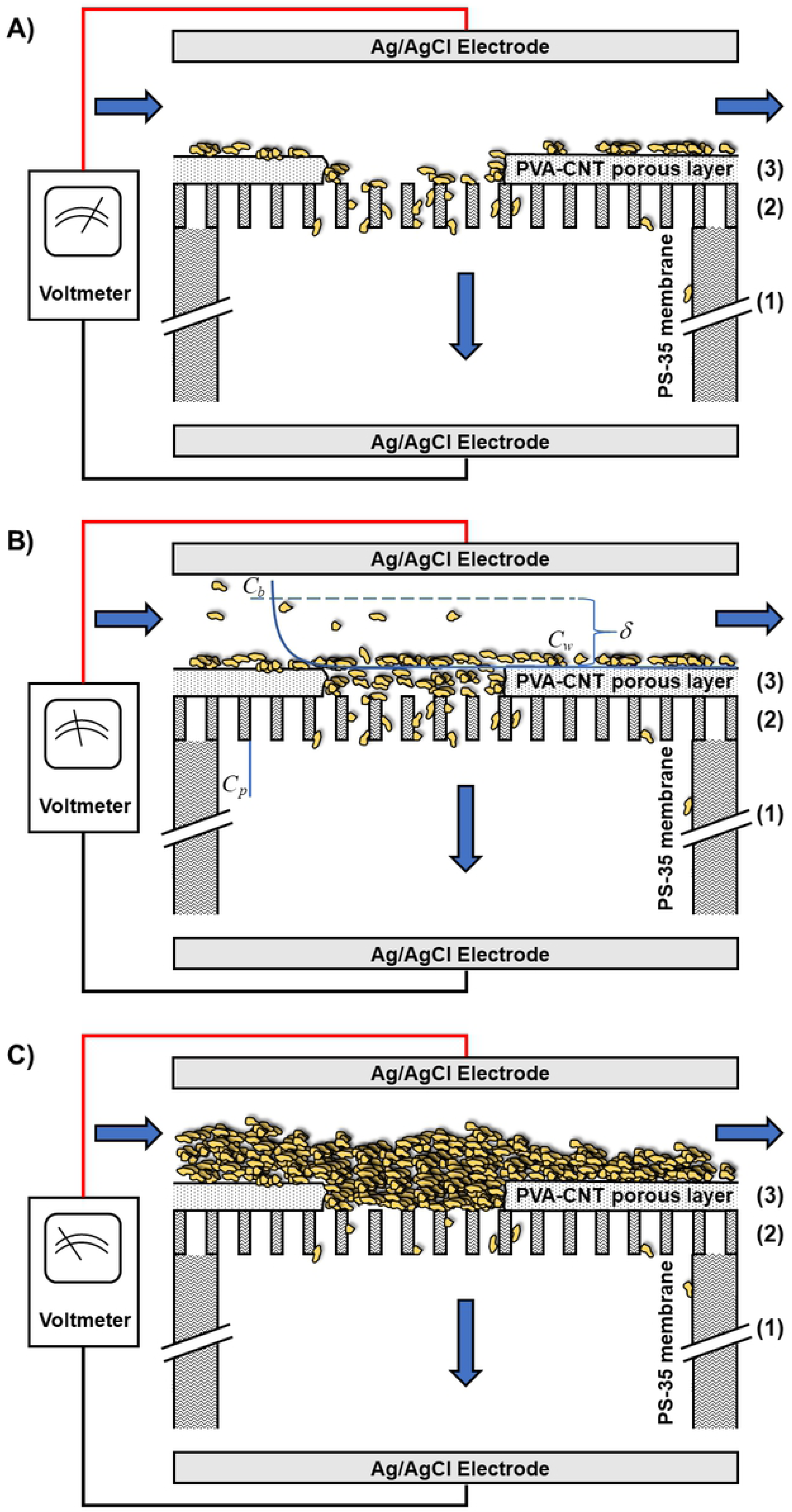
PVA-CNT layer makes a significant contribution to the global zeta potential due to fouling at the thin film surface. Schematic of streaming potential setup for measurement of the global zeta potential across the membrane pore walls of the (1) PS-35 support layer, (2) PS-35 skin layer, and (3) PVA-CNT layer (not to scale) under three mechanisms of fouling and concentration polarization above the surface of the PVA-CNT layer: A) minimal fouling and negligible additional hydraulic resistance due to concentration polarization, B) significant concentration polarization, C) extensive fouling. Under A), the PS-35 support layer dominates the contribution to the global zeta potential relative to the other layers due to the significant pressure drop across the thick support layer. Under B), while proteins transported through the membrane give rise to filtrate with a specific permeate concentration (C_p_), proteins retained by the PVA-CNT layer result in an elevated wall concentration (C_w_) relative to the bulk concentration (C_b_). Concentration polarization across the boundary layer (δ) results in significant contributions to the global zeta potential due to the additional hydraulic resistance. Conversely, under C), the PVA-CNT layer fouled with adsorbed protein plays a significant role in the contribution to the global zeta potential due to the reduced effective porosity and pore size and increased layer thickness.

For single protein UF/EUF of 0.1 g/L HEL, the evaluated zeta potentials of the PS-35 membrane after UF at 0 V (vs. Ag/AgCl), the PVA-CNT membrane after UF at 0 V, and the PVA-CNT membrane after EUF at −4.6 V were −3.9 ± 0.1 mV, −15.9 ± 0.2 mV, and −3.0 ± 0.1 mV, respectively (Table 2). The magnitude of the negative zeta potential of the PS-35 membrane following single protein UF of HEL at 0 V vs. Ag/AgCl was significantly less than that of the virgin PS-35 membrane due to substantial fouling of HEL within the polysulfone membrane pore walls which modifies the surface net charge to more closely reflect that of the protein. The fouling may be attributed to the electrostatic attraction of the positively charged HEL (at the system pH of 7.4) to the negatively charged PS-35 membrane as well as the hydrophobic interactions between the protein and membrane. For the composite PVA-CNT membrane, the nominal value of the zeta potential was only slightly lower than that of the virgin membrane. This suggests that HEL proteins were adsorbed onto the thin PVA-CNT layer. Interestingly, upon application of an electric potential of −4.6 V (vs. Ag/AgCl) during single protein EUF of HEL using the PVA-CNT membrane, the nominal value of the zeta potential was significantly reduced. Although the PVA-CNT layer is relatively thin and has a relatively large pore size, it is possible that concentration polarization above the interface resulted in significant additional hydraulic resistance and subsequent reduction in the measured global zeta potential (Fig 6B). Alternatively, the significant reduction in the global zeta potential could be explained by fouling on the PVA-CNT layer. As discussed earlier, the thick support layer of the polysulfone membrane should dominate the contribution to the measured global zeta potential over the thin PVA-CNT network and the selective layer. However, the contribution of the PVA-CNT layer to the global zeta potential increases with decreasing pore radius ratios (*a^(1)^*/*a^(3)^*) and length fractions of the support layer (*X*) [49]. The more dramatic reduction in the nominal global zeta potential of the PVA-CNT composite membrane upon application of the electric potential could therefore be explained by heavy fouling on the surface of the PVA-CNT layer resulting from the larger electrostatic forces of attraction between the positively charged HEL proteins and the negatively charged PVA-CNT network (Fig 6C). The significant fouling during single protein EUF of HEL results in a dramatic decrease in the apparent pore radius of the PVA-CNT network.

For single protein UF/EUF of 0.1 g/L αLA, the evaluated zeta potentials of the PS-35 membrane after UF at 0 V (vs. Ag/AgCl), the PVA-CNT membrane after UF at 0 V, and the PVA-CNT membrane after EUF at −4.6 V were −0.3 ± 0.1 mV, −4.3 ± 0.1 mV, and −9.9 ± 0.1 mV, respectively. The significantly reduced nominal zeta potential of the fouled PS-35 membrane following single protein UF of αLA compared to that of the virgin membrane suggests the proteins are adsorbed along the PS-35 membrane walls and the PVA-CNT network. The PS-35 membrane fouled with a single protein solution of HEL exhibited a more negative zeta potential compared to the PS-35 membrane fouled with a single protein of αLA which may be due to the HEL proteins being adsorbed in a configuration that exposes their negative sites to the outer side of the adsorbed layer [52]. The PS-35 membrane with the added PVA-CNT layer exhibited a larger nominal zeta potential over the PS-35 only counterpart following single protein EUF of αLA due to electrostatic repulsion of the proteins and subsequent reduced fouling within the PS-35 membrane pores. A further increase in the nominal zeta potential was observed upon application of the cathodic potential during single protein EUF of αLA due to the larger electrostatic repulsive force.

For binary protein UF/EUF of feed solution containing 0.1 g/L HEL and 0.1 g/L αLA, the evaluated zeta potentials of the PS-35 membrane after UF at 0 V (vs. Ag/AgCl), the PVA-CNT membrane after UF at 0 V, and the PVA-CNT membrane after EUF at −4.6 V were 0.8 ± 0.1 mV, −5.7 ± 0.1 mV, 1.5 ± 0.1 mV, respectively. The positive zeta potential of the PS-35 membrane following binary protein EUF of HEL and αLA indicates significant fouling along the membrane pores. Compared to the virgin PVA-CNT membrane, the PVA-CNT membrane post-experiment at no applied electric potential exhibited a reduced nominal zeta potential. Although this suggests there was still fouling along the pore walls of the bottom PS-35 membrane, the fouling along the polysulfone pores of the composite membrane is not as significant compared to that of the PS-35 only membrane due to protein adsorption within the PVA-CNT network. Application of an electric potential of −4.6 V (vs. Ag/AgCl) during binary protein EUF using the PVA-CNT membrane resulted in a positive zeta potential which suggests heavy protein adsorption and aggregation above and within the PVA-CNT layer.

Visualizations of the surfaces of the PVA-CNT membranes following an extended duration of EUF (feed: 0.1 g/L of each protein component, 4 mM ionic strength, pH 7.4; operating conditions: 555 s^-1^ crossflow shear rate, 1 psi TMP, 9.33 h) were obtained by SEM and are displayed in Fig 7. Additional SEM images of the PS-35 membranes post-filtration are provided in the Supporting Information (S9 Fig). While the global zeta potential values calculated from streaming potential measurements provide information regarding adsorption within both the PVA-CNT porous network and the polysulfone membrane pores, scanning electron microscopy gives insight to fouling at the upper surface of the PVA-CNT film.

**Fig 7.**
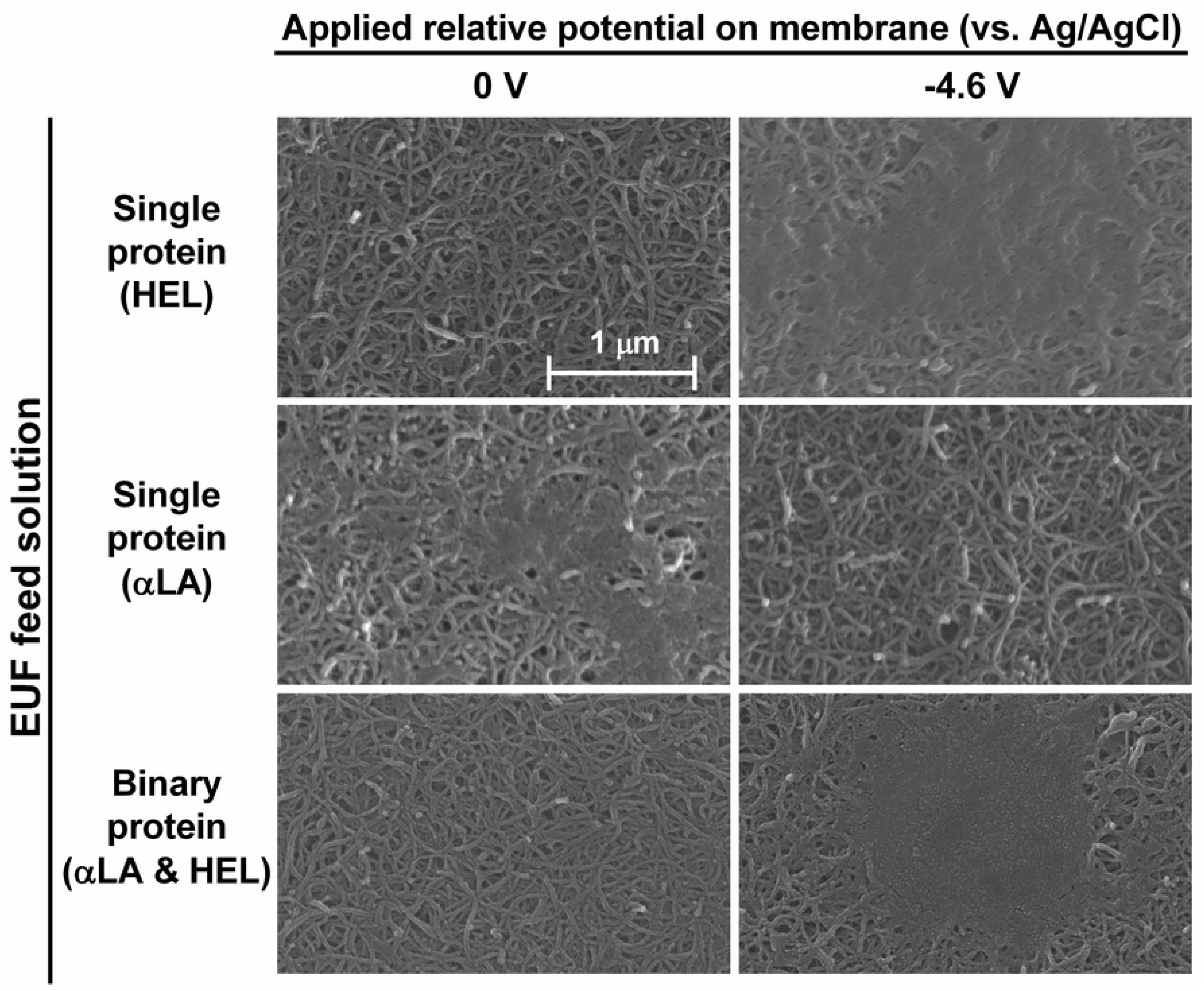
Protein adsorption and aggregation develop above the PVA-CNT membrane surfaces following crossflow EUF. Fouling above the PVA-CNT membrane surfaces following protein crossflow EUF (0.1 g/L of each protein component; 555 s-1 crossflow shear rate; 1 psi TMP; 9.33 h duration; and 0 V and −4.6 V potentials (vs. Ag/AgCl) applied to the PVA-CNT membrane). Images were obtained by SEM. Application of an external electric potential results in increased fouling for single protein HEL EUF and decreased fouling for single protein αLA EUF due to electrostatic interactions between the charged protein species and the negatively charged PVA-CNT layer. Application of an electric potential during binary protein EUF of αLA and HEL results in increased fouling due to a combination of protein-protein and protein-PVA-CNT layer electrostatic interactions. The observed multilayer protein adsorption leads to pore narrowing and clogging.

At the experimental pH of 7.4, large complexes of protein deposits on the top surface of the PVA-CNT membranes were observed following single protein HEL and binary protein (HEL and αLA) EUF as a result of electrostatic attraction between the positively charged HEL and the PVA-CNT layer, and additionally for the binary protein case, a result of protein-protein interactions leading to heavy aggregation and adsorption. Conversely, reduced surface fouling was observed when applying a cathodic potential during EUF of αLA due to the electrostatic repulsion of the negatively charged proteins from the membrane. The visualizations of fouling at the surface of the PVA-CNT layer suggest heavy adsorption significantly affected the measured global zeta potentials. Although the observed protein fouling above the PVA-CNT membrane following EUF was different in the cases of no applied potential and −4.6 V potential (vs. Ag/AgCl), it is worth noting that the observed protein sieving for EUF with an applied potential still converged to that without an applied potential at extended durations of EUF. This suggests protein transmission at the longer time periods of EUF is still predominantly influenced by the protein adsorption within the electrically conductive network and membrane pores. The SEM images of the surfaces post-EUF, along with the permeate flux, sieving, and zeta potential measurements, confirm that protein adsorption and aggregation are significant factors in protein EUF with an electrically conductive membrane.

## Conclusions

Electroultrafiltration using a CNT-based composite ultrafiltration membrane was evaluated for its effectiveness in improving membrane performance during crossflow filtration of dilute single and binary protein solutions of α-lactalbumin and hen egg-white lysozyme (0.1 g/L of each species; pH 7.4; 4 mM ionic strength; 1 psi TMP). The effects of the presence of the PVA-CNT layer and the applied electric potentials on the transient permeate flux, protein sieving, and protein selectivity were investigated. Non-ideality in the permeate flux behavior characterized by abrupt reductions in normalized permeate flux were observed in single protein studies for αLA and HEL. A significant reduction in the normalized permeate flux was observed for binary protein studies with and without an applied cathodic potential. Application of a cathodic potential of −4.6 V (vs. Ag/AgCl) across the PVA-CNT membrane yielded a temporary enhancement in protein sieving for both lysozyme and α-lactalbumin and in protein selectivity (*S_o,αLA_ / S_o,HEL_*).

The timepoints of the inflection in protein sieving are consistent with the timepoints corresponding to the dramatic reductions in permeate flux suggesting electrostatic interactions between the fouled protein layer and adjacent proteins dominate the protein transport behavior in later stages of EUF. Characterization of membrane fouling through measurements of the transient permeate flux, as well as zeta potential measurements and SEM visualizations of the conditioned membranes, demonstrated increased protein adsorption with an applied electric potential. Through consideration of the relative contributions of each layer of the three-layer composite PVA-CNT membrane along with coupled analysis of the global zeta potential and the surface fouling shown by SEM, the reduction in the steady-state permeate flux and protein sieving during protein EUF was determined to be due to significant fouling above and within the PVA-CNT network. Protein transport through the electrically conductive PVA-CNT membranes at extended periods of EUF is ultimately dependent on the electrostatic interactions between the unbound proteins and adsorbed proteins on the pore walls of the PVA-CNT network and skin layer of the PS-35 membrane.

While it was not possible to maintain the enhancement in protein sieving during EUF with a cathodic membrane over extended periods of treatment due to protein adsorption, the EUF process may have potential as an alternate mode of operation for specific protein separation processes including preferential filtration of similarly sized charged proteins. The PVA-CNT layer may also be utilized as a replaceable adsorbent layer for cascade ultrafiltration systems [53]. The tunability of protein transport through the CNT-polymer composite membrane over a short time period may also be utilized in drug delivery applications. Protein binding to the multiwalled carbon nanotubes may be controlled by changing the functionalization and nanotube diameter to modulate the protein transport behavior appropriate for various applications [19, 21].

## Acknowledgments

This research was funded by an endowment from Jacques S. Yeager, Sr. Professorship. We would like to thank Davisco Foods International, Inc. (Agropur, Inc.) for providing the α-lactalbumin used in the study. Special thanks to Dr. Alexander Dudchenko for providing valuable feedback in our discussion

## Supporting Information

**S8 Fig. Digital camera image of PVA-CNT PS-35 composite membrane.**

**S9 Fig. SEM image of cross section of CNT layer deposited on a polysulfone UF membrane.**

**S10 Fig. Evaluation of PVA-CNT thin film thickness from SEM image of cross section of CNT layer.**

**S11 Fig. SEM image of top view of a virgin PVA-CNT membrane at A) 50K magnification and B) 5K magnification.**

**S12 Fig. Pore size analysis of PVA-CNT using ImageJ.** A) Original PVA-CNT membrane surface image from SEM; B) Binary image from thresholding; C) Output of analyzed regions used for calculation of pore size.

**S13 Fig. Schematic of experimental flow cell.** A) Top view of membrane in flow cell. The potential was monitored using a multimeter at each side of the membrane at locations adjacent to the edge of the flow cell and the edge of the membrane. B) Side view of the membrane flow cell. A plastic mesh spacer was placed between the PVA-CNT membrane and the carbon mesh electrode.

**S14 Fig. Charge distribution of hen egg-white lysozyme and α-lactalbumin.** Electrostatic potential visualization of A) hen egg-white lysozyme (PDB ID: 1HFZ) and B) α-lactalbumin (PDB ID: 1HFZ) from calculations using the APBS-PDB2PQR software suite and generated using VMD. The potentials are on a [-4 k_B_T/e, 4 k**_B_**T/E] red-white-blue color map. Calculations were performed at 4 mM ionic strength, 298.15 Kelvin, protein dielectric of 2, and solvent dielectric of 78. Although α-lactalbumin has a net negative charge, the protein contains positively charged regions which may bind to the negatively charged CNTs and the polysulfone membrane.

**S15 Fig. Binary protein electroultrafiltration with application of anodic and cathodic potentials.** A) Normalized permeate flux, B) observed sieving coefficient, and C) mass flux during EUF of binary protein solutions of 0.1 g/L αLA and 0.1 g/L HEL at TMP of 1 psi with different cathodic potentials (vs. Ag/AgCl) applied to the PS-35 (n=3) and PVA-CNT membranes (n=2; n=1 for 2.5 V). Error bars represent the standard error of the weighted mean.

**S3 Table. Hydraulic permeability for assessment of membrane fouling.** Weighted average pre-experimental and post-experimental hydraulic permeability data of membranes used in EUF experiments (1 psi, 555 s-1 crossflow shear rate, 4 mM ionic strength, pH 7.4).

**S16 Fig. SEM image of surfaces of uncoated PS-35 polymeric ultrafiltration membranes.** A) virgin PS-35 membrane, B) PS-35 membrane following single protein HEL UF (0 V vs. Ag/AgCl), C) PS-35 membrane following single protein αLA UF (0 V vs. Ag/AgCl), D) binary protein UF of αLA and HEL(0 V vs. Ag/AgCl) with 0.1 g/L of each protein component; 555 s-1 crossflow shear rate; 1 psi TMP; 9.33 h duration.

**S4 Table. Debye length calculations at low and high surface potentials.** Although the calculated Debye lengths are significantly smaller than the large pore sizes of the PVA-CNT network, the fouling observed on the surface extend across the porous PVA-CNT thin film which suggests formation of multilayers of adsorbed protein.

